# SARS-CoV-2 Variants are Selecting for Spike Protein Mutations that Increase Protein Stability

**DOI:** 10.1101/2021.06.25.449882

**Authors:** David Shorthouse, Benjamin. A. Hall

**Affiliations:** Department of Medical Physics and Biomedical Engineering, University College London, Gower Street, London, WC1E 6BT

**Keywords:** COVID-19, SARS-CoV-2, Spike protein, variants, mutation, stability, selection

## Abstract

The emergence of SARS-CoV-2 in 2019 has caused severe disruption and a huge number of human deaths across the globe. As the pandemic spreads, a natural result is the emergence of variants with a variety of amino acid mutations. Variants of SARS-CoV-2 with mutations in their spike protein may result in an increased infectivity, increased lethality, or immune escape, and whilst many of these properties can be explained through changes to binding affinity or changes to post-translational modification, many mutations have no known biophysical impact on the structure of protein. The Gibbs free energy of a protein represents a measure of protein stability, with an increased stability resulting in a protein that is more thermodynamically stable, and more robust to changes in external environment.

Here we show that mutations in the spike proteins of SARS-CoV-2 are selecting for amino acid changes that result in a more stable protein than expected by chance. We calculate all possible mutations in the SARS-CoV-2 spike protein, and show that many variants are more stable than expected when compared to the background, indicating that protein stability is an important consideration for the understanding of SARS-CoV-2 evolution. Variants exhibit a range of stabilities, and we further suggest that some stabilising mutations may be acting as a “counterbalance” to destabilising mutations that have other properties, such as increasing binding site affinity for the human ACE2 receptor. We suggest that protein folding calculations offer a useful tool for early identification of advantageous mutations.

---

TEXT: Since the emergence of SARS-CoV-2 in late 2019, over 2 million people have died as a result of infection^1^. As the global pandemic continues, the emergence of viral variants with RNA mutations is an expected phenomena, caused by random errors in RNA copying, and selected for by evolutionary pressure^2^. Many variants contain mutations in the spike protein that confer an advantage to the virus, such as increased ACE2 receptor binding^3^, glycosylation/cleavage site alterations^4^, and immune evasion^5^, as well as protein stability^6^. Understanding these properties helps infer how a variant may differ from another mutational profile, and provides insights into the mechanisms by which variants differ, such as increased infectivity or vaccine resistance^7–9^. The WHO classifies variants in SARS-CoV-2 into major categories, the two most important: “Variants of Concern” and “Variants of Interest” are assigned to emerging variants likely to have a different phenotype and mutational profile to the original SARS-CoV-2^10^.

Changes in Gibbs Free Energy (called ΔΔG) is a measure of the thermodynamic energy change upon amino acid mutation in a protein. Prediction of the changes in ΔΔG are routinely used in protein engineering and design for optimization of enzymes or stabilisation of protein complexes^11,12^, and we have recently shown that they can be predictive of mutations that destabilise or damage a protein in a cancer context^13–15^. Whilst stability of mutations has been assessed in the SARS-CoV-2 spike protein^16,17^, variant analysis has not yet been performed. Mutations that stabilise the SARS-CoV-2 spike protein are likely to lead to better binding to other molecules, and a greater lifespan of a protein before thermal unfolding. The requirement for calculation of predicted ΔΔG values is protein structural information, which was recently published for the SARS-CoV-2 spike protein^18^.

Here we calculate the ΔΔG on mutation for every possible missense mutation in the SARS-CoV-2 spike protein. With this “background” mutation rate we show that mutations to the spike protein observed in emerging SARS-CoV-2 variants have a lower ΔΔG, and a higher proportion of stabilising mutations than expected. We further show that combinations of mutations result in synergistic stability changes, and so highlight possible evolutionary orders of mutations. This suggests an important role for protein stability when considering the evolution of SARS-CoV-2.

The SARS-CoV-2 spike protein is composed of a trimer of 3 identical subunits (Figure 1a) that sits in the membrane of the virion and interacts with the human ACE2 receptor to facilitate infection of a host cell. The structure of the spike protein was recently elucidated, enabling the calculation of predictive ΔΔG values for mutations. The “Alpha” variant, first identified in December 2020 in the United Kingdom^19,20^ has been found to be more transmissible than the original virus, with an increased affinity for binding the human ACE2 receptor^21^, and by April 2020 had become the most dominant variant in the UK. The Alpha variant carries 23 common mutations across its genome, 7 of which are amino acid substitution mutations in the spike protein, 6 of which are at locations for which crystallographic data is available (Figure 1b).

**Figure 1.**
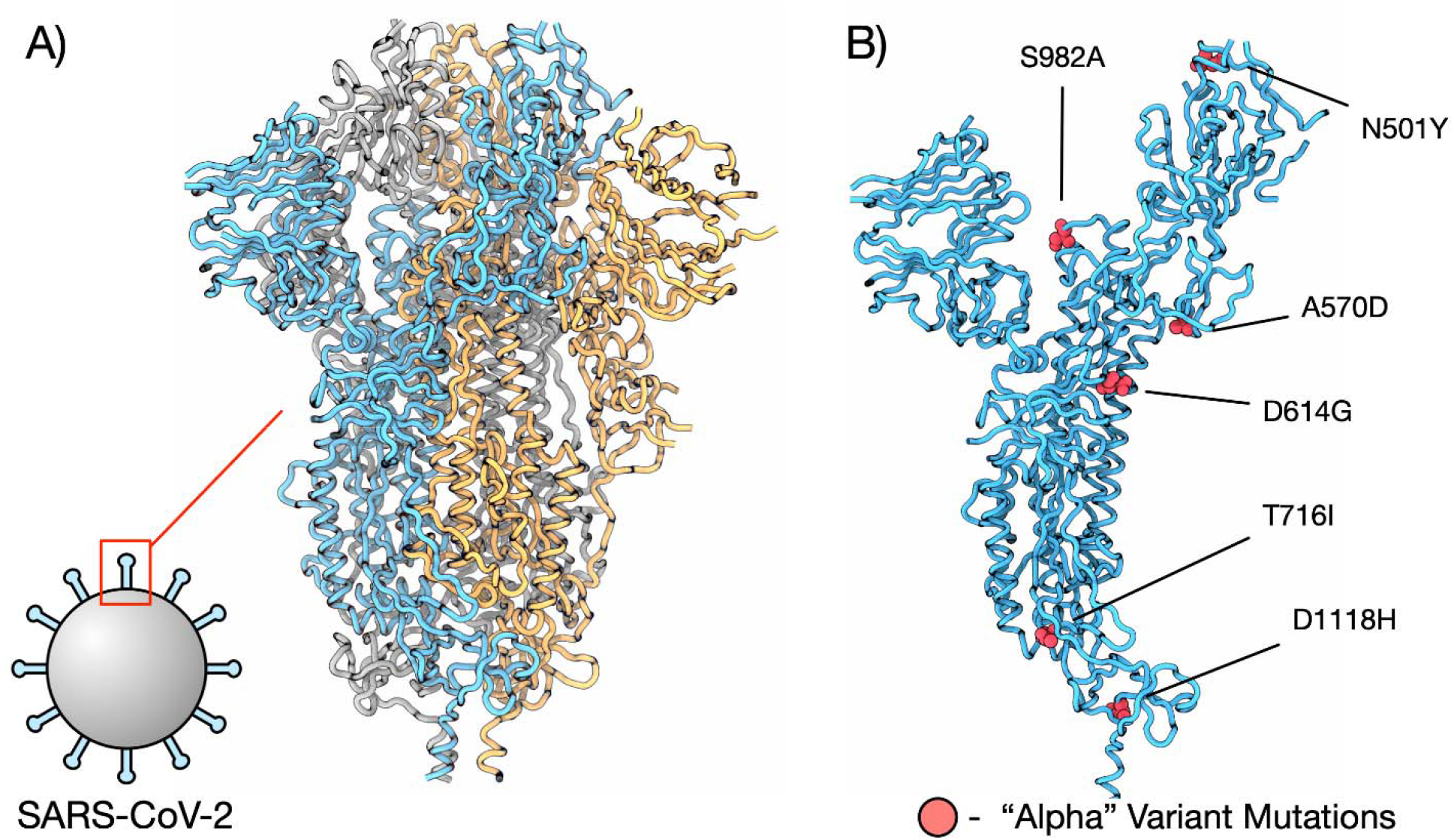
Structure of SARS-CoV-2 spike protein. A) Full structure of COVID spike protein structure (PDBID 6XVV), subunits are coloured blue, orange, and grey. B) Single subunit of spike protein with residues from the “Alpha” variant coloured red and labelled.

To explore the mutational landscape of the spike protein and SARS-CoV-2 variants, we first calculated the ΔΔG for each of the possible 19440 missense mutations in the 6XVV cryo-em structure using FoldX^22^ (Supplementary Table 1). As is consistent with previous studies, we find mutations that stabilise the protein are rare (Figure 2a). We define mutations with an induced ΔΔG of < -1 Kcal/mol as strongly stabilising, and mutations with a ΔΔG > 2.5 Kcal/mol as strongly destabilising, with those between zero and each threshold described as mildy stabilising and destabilising respectively. Only 767 (3.9%) of possible mutations are predicted to strongly stabilise the protein, and only 3699 (19%) have a ΔΔG < 0. With this “background” mutational distribution, we compared to mutations found only in WHO “Variants of Concern” and “Variants of Interest” as of June 2021 (Figure 2b). Mutations found in both categories have a significantly lower ΔΔG (t-test pvalue < 0.05) than bulk population, indicating that variants may be evolutionarily selecting for stabilising or non-destabilising mutations. Considering individual mutations found in “Variants of Concern” (Figure 2c), none of the mutations observed induce a ΔΔG > 2.5 Kcal/mol (defined as strongly destabilising), significantly different to the expected 34% of possible mutations meeting this threshold (chi squared pvalue <0.05). Additionally 4 of 17 mutations have ΔΔG of <= ∼-1 Kcal/mol (N501Y, H655Y, T716I, and T1027I), showing a significant enrichment for stabilising mutations in the “Variants of Concern” (chi squared pvalue <0.05). We conclude that there is a statistical enrichment for mutations that stabilise the spike protein compared to the mutational background in SARS-CoV-2 variants.

**Figure 2.**
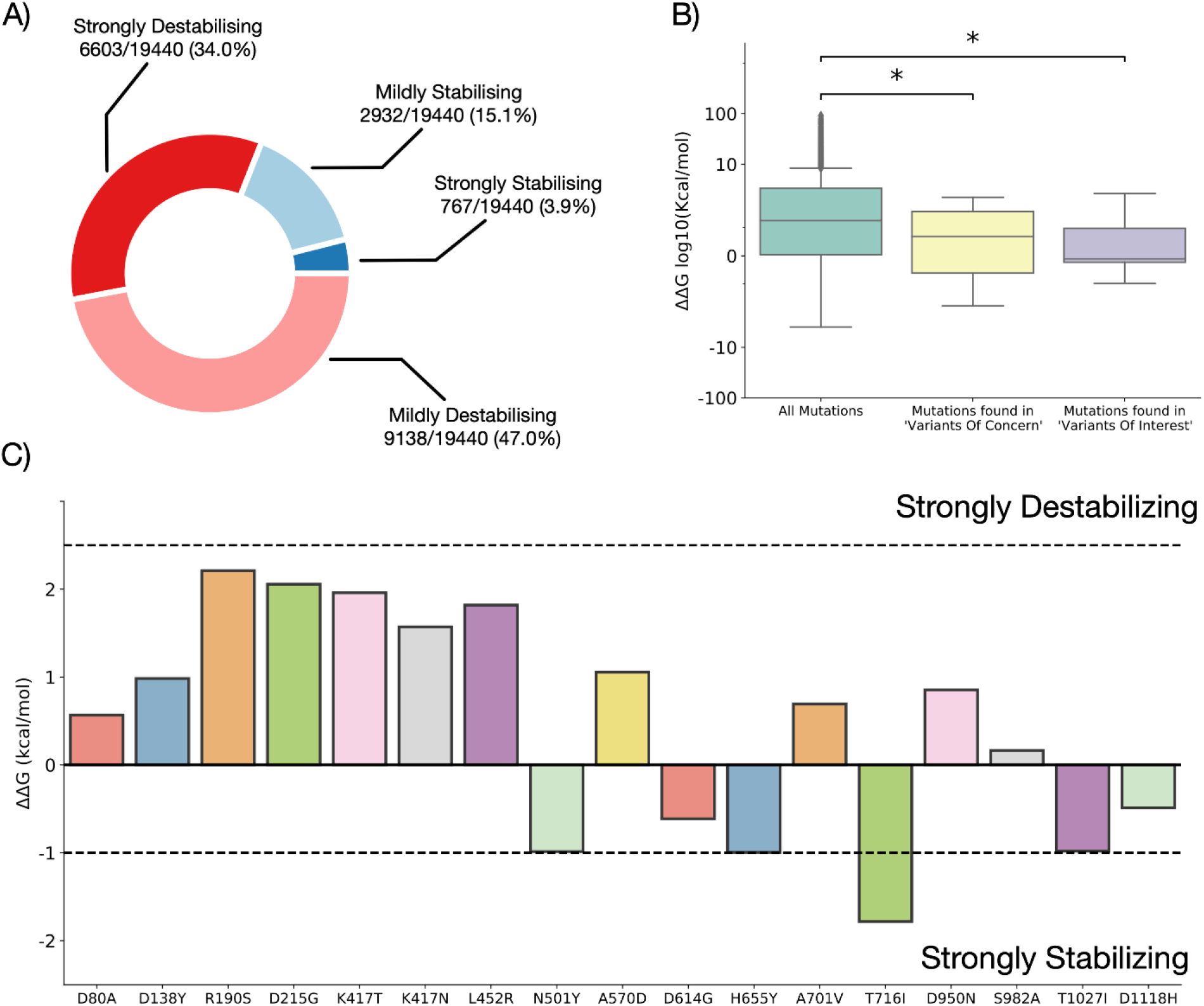
Saturation screen of SARS-CoV-2 spike protein. A) Proportion of mutations that are strongly stabilising (ΔΔG < -1 Kcal/mol), strongly destabilising (ΔΔG > 2.5 Kcal/mol), mildly stabilising (0 > ΔΔG < -1 Kcal/mol, or mildly destabilising (0 < ΔΔG < 2.5 Kcal/mol). B) ΔΔG for all 19440 mutations compared to those in WHO “Variants of Concern” and “Variants of Interest”. * represents ttest pvalue <= 0.05. C) ΔΔG (Kcal/mol) for all mutations observed in WHO “Variants of Concern”.

To further unpick the stability of specific SARS-CoV-2 variants we calculated the ΔΔG distribution for mutations individually for each variant in the two categories (Figure 3, variants and mutations included in Table 1). Of the 10 variants studied, 7 have a statistically significantly lower ΔΔG than the bulk mutational background (ttest pvalue <=0.05), and 5 variants (Alpha, Gamma, Eta, Theta, and Iota) have a mean ΔΔG less than 0, indicating that the protein will be stabilised with respect to the original variant. To further study this relationship we calculated the expected ΔΔG for each variant given the number of mutations occurring in it (Supplementary Figure 1), and find that all variants aside from Beta and Eta have a lower ΔΔG than expected given the number of mutations they contain. Of particular note is the Alpha variant, which has a ΔΔG distribution almost entirely below zero, as well as the emerging Theta variant first identified in India.

**Figure 3.**
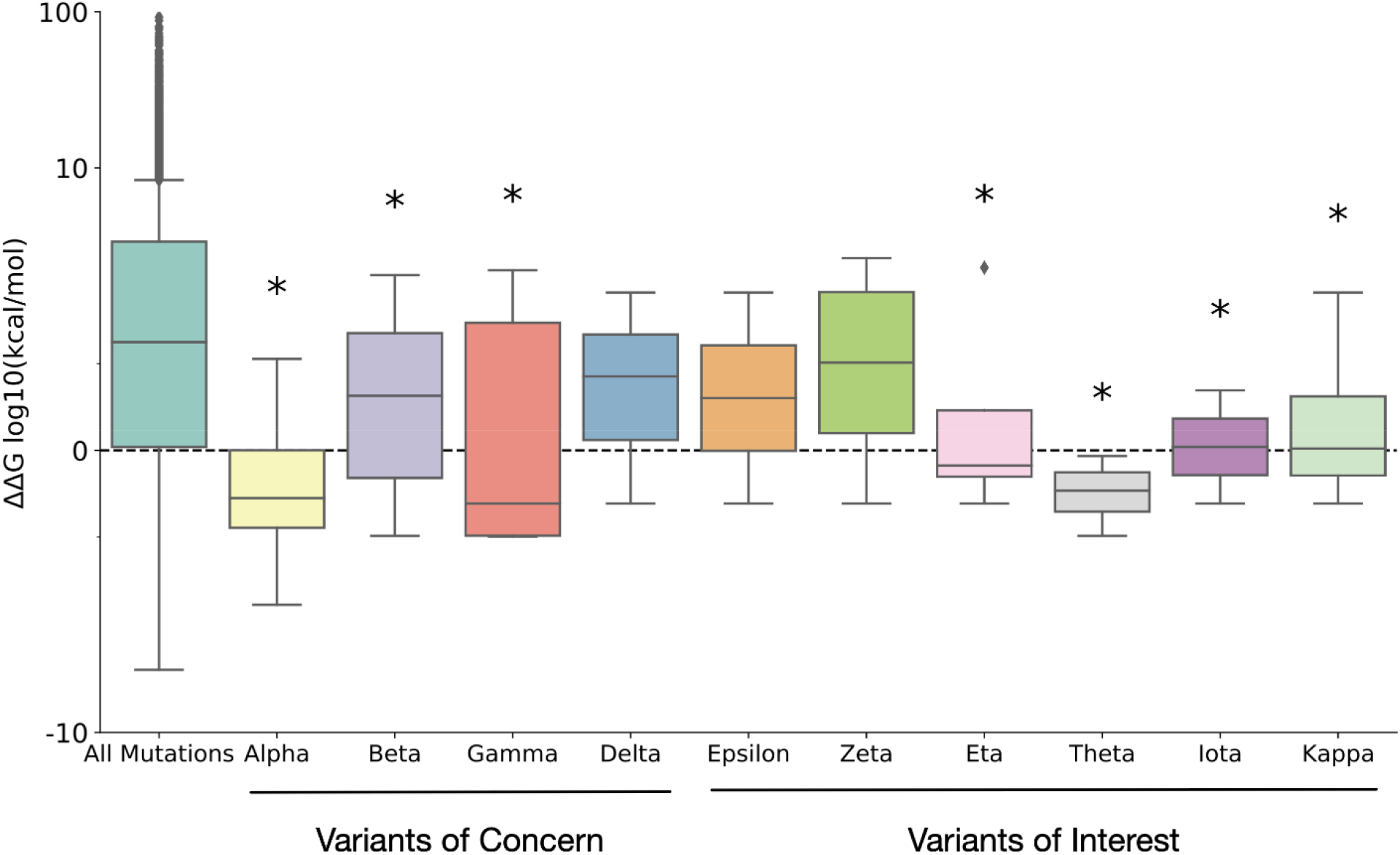
ΔΔG (Kcal/mol) for all mutations observed in WHO “Variants of Concern” and “Variants of Interest” compared to all 19440 possible mutations in the SARS-CoV-2 spike protein. * represents ttest pvalue <=0.05.

**Table 1:** SARS-CoV-2 Variants of Concern (Alpha, Beta, Gamma, and Delta), and Variants of Interest (Epsilon, Zeta, Eta, Theta, Iota, and Kappa) as of June 2021. * indicates a mutated residue is not included in the 6XVV structure.

| WHO Label | PANGO Lineage | Location Identified | Mutations Present |
| --- | --- | --- | --- |
| Alpha | B.1.1.7 | United Kingdom | N501Y, A570D, D614G, P681H*, T716I, S982A, D1118H |
| Beta | B.1.351 | South Africa | D80A, D215G, K417N, E484K*, N501Y, D614G, A701V |
| Gamma | P.1 | Brazil | L18F*, T20N*, P26S*, D138Y, R190S, K417T, E484K*, N501Y, D614G, H655Y, T1027I, V1176F* |
| Delta | B.1.617.2 | India | T19R*, R158G*, L452R, T478K*, D614G, P681R*, D950N |
| Epsilon | B.1.427 | United States | S13I*, W152C*, L452R, D614G |
| Zeta | P.2 | Brazil | E484K*, F565L, D614G, V1176F* |
| Eta | B.1.525 | Multiple Countries | Q52R, A67V, E484K*, D614G, Q677H*, F888L |
| Theta | P.3 | Philippines | E484K*, N501Y, D614G, P681H*, E1092K, H1101Y, |
|  |  |  | V1176F* |
| Iota | B.1.526 | United States | L5F*, T95I*, D253G*, E484K*,<br>D614G, A701V |
| Kappa | B.1.617.1 | India | G142D, E154K*, L452R,<br>E484Q*, D614G, P681R*,<br>Q1071H |

Finally, to study the potential evolutionary order and gain insights into mechanisms of mutational selection, we calculated the ΔΔG for every possible combination of mutations in each variant, shown in Figure 4 for the Alpha variant (Supplementary Figures 2-9 for other variants). For the alpha variant there is a consistent trend towards stabilisation as more mutational combinations are considered, with all combinations of 5 or more mutations (of the 6 possible to model in the structure) resulting in a predicted stabilisation of the protein with respect to the original. Furthermore we observe combinations that result in a positive ΔΔG, which are likely to be evolutionarily less favourable (when considering stability alone), and so we expect that these combinations would be less likely to occur in the evolutionary history than stabilising combinations. Furthermore, some variants, such as the Beta variant first identified in South Africa in May 2020, contains combinations of mutations with a ΔΔG expected to be highly destabilising, and whilst the final ΔΔG of all mutations is still predicted to be strongly destabilising, it is reduced compared to the most extreme combinations, indicating that a potential driver of selection of other mutants may be that they stabilise the protein complex enough for it to function, whilst retaining the advantageous properties unrelated to stability from the destabilising mutations.

**Figure 4.**
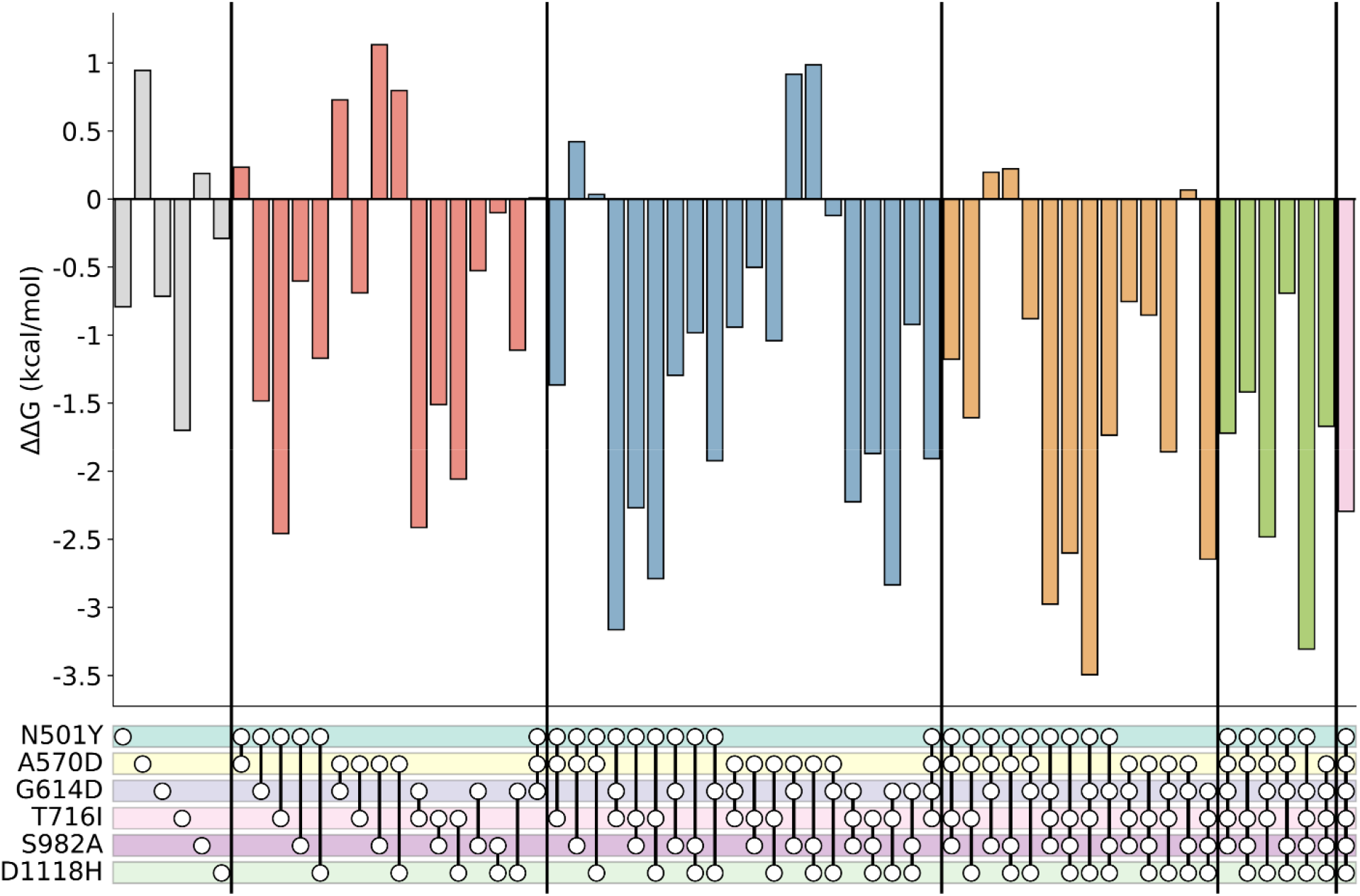
Mutational combinations in the Alpha variant. Upset plot of ΔΔG (Kcal/mol) for all combinations of 6 mutations in the Alpha variant. Dot and line represents presence of connected mutations in the calculation.

This work highlights that mutations with a stabilising effect on the SARS-CoV-2 spike protein are one of the key drivers of evolution of the virus, and contributing to the increased transmissibility of emerging variants. That variants are more stable than expected by chance shows that evolution is favouring mutations with a stabilising effect, and it may be that mutations that destabilise a protein but have other influences on protein structure, such as K417N^21^, which alters ACE2 binding affinity, are offset or preceded by mutations that stabilise the structure. We note however, that not all mutations in all variants can be considered, due to missing regions of the cryo-em structure, and as such this study does not necessarily represent the true ΔΔG for each variant. Furthermore, we study only the structure in its “closed” conformation as we feel this is the most physiological relevant of the existing structures, but further work will need to address the impact of the dynamic range of the structure on mutational stability. We highlight that stability of the SARS-CoV-2 spike protein is an important consideration for future study of variants, and is likely one of a number of driving forces in the evolution of new variants. Finally, we suggest that folding simulations of newly sequenced variants may offer a computationally inexpensive method to highlight advantageous mutations, and that prospective simulation of further mutation to these samples may predict future variants to surveil for.

## ASSOCIATED CONTENT

### Supporting Information

The following files are available free of charge.

### Supplementary Information

Methods for ΔΔG calculation and supplementary figures (PDF)

**Supplementary Table 1**: Table containing predicted ΔΔG for every possible mutations in SARS-CoV-2 structure PDBID 6VXX (XLSX)

### Author Contributions

DS and BAH conceived the study and wrote the manuscript. DS generated all data and performed all analysis.

### Funding Sources

This work was supported by the Medical Research Council (grant no. MR/S000216/1). B.A.H. acknowledges support from the Royal Society (grant no. UF130039). The authors declare no competing financial interest.

## REFERENCES

(1) Peacock, T. P.; Penrice-Randal, R.; Hiscox, J. A.; Barclay, W. S. SARS-CoV-2 One Year on: Evidence for Ongoing Viral Adaptation. Journal of General Virology 2021, 102 (4).

(2) Plante, J. A.; Mitchell, B. M.; Plante, K. S.; Debbink, K.; Weaver, S. C.; Menachery, V. D. The Variant Gambit: COVID-19’s next Move. Cell Host & Microbe 2021, 29 (4), 508–515.

(3) Ozono, S.; Zhang, Y.; Ode, H.; Sano, K.; Tan, T. S.; Imai, K.; Miyoshi, K.; Kishigami, S.; Ueno, T.; Iwatani, Y.; Suzuki, T.; Tokunaga, K. SARS-CoV-2 D614G Spike Mutation Increases Entry Efficiency with Enhanced ACE2-Binding Affinity. Nature Communications 2021, 12 (1), 848.

(4) Tortorici, M. A.; Veesler, D. Structural Insights into Coronavirus Entry. In Advances in Virus Research; 2019; Vol. 105, pp 93–116.

(5) Harvey, W. T.; Carabelli, A. M.; Jackson, B.; Gupta, R. K.; Thomson, E. C.; Harrison, E. M.; Ludden, C.; Reeve, R.; Rambaut, A.; Peacock, S. J.; Robertson, D. L. SARS-CoV-2 Variants, Spike Mutations and Immune Escape. Nature Reviews Microbiology 2021, 19 (7), 409–424.

(6) Berger, I.; Schaffitzel, C. The SARS-CoV-2 Spike Protein: Balancing Stability and Infectivity. Cell Research 2020, 30 (12), 1059–1060.

(7) Planas, D.; Bruel, T.; Grzelak, L.; Guivel-Benhassine, F.; Staropoli, I.; Porrot, F.; Planchais, C.; Buchrieser, J.; Rajah, M. M.; Bishop, E.; Albert, M.; Donati, F.; Prot, M.; Behillil, S.; Enouf, V.; Maquart, M.; Smati-Lafarge, M.; Varon, E.; Schortgen, F.; Yahyaoui, L.; Gonzalez, M.; de Sèze, J.; Péré, H.; Veyer, D.; Sève, A.; Simon-Lorière, E.; Fafi-Kremer, S.; Stefic, K.; Mouquet, H.; Hocqueloux, L.; van der Werf, S.; Prazuck, T.; Schwartz, O. Sensitivity of Infectious SARS-CoV-2 B.1.1.7 and B.1.351 Variants to Neutralizing Antibodies. Nature Medicine 2021, 27 (5), 917–924.

(8) Korber, B.; Fischer, W. M.; Gnanakaran, S.; Yoon, H.; Theiler, J.; Abfalterer, W.; Hengartner, N.; Giorgi, E. E.; Bhattacharya, T.; Foley, B.; Hastie, K. M.; Parker, M. D.; Partridge, D. G.; Evans, C. M.; Freeman, T. M.; de Silva, T. I.; McDanal, C.; Perez, L. G.; Tang, H.; Moon-Walker, A.; Whelan, S. P.; LaBranche, C. C.; Saphire, E. O.; Montefiori, D. C.; Angyal, A.; Brown, R. L.; Carrilero, L.; Green, L. R.; Groves, D. C.; Johnson, K. J.; Keeley, A. J.; Lindsey, B. B.; Parsons, P. J.; Raza, M.; Rowland-Jones, S.; Smith, N.; Tucker, R. M.; Wang, D.; Wyles, M. D. Tracking Changes in SARS-CoV-2 Spike: Evidence That D614G Increases Infectivity of the COVID-19 Virus. Cell 2020, 182 (4), 812–827.e19.

(9) Khan, A.; Zia, T.; Suleman, M.; Khan, T.; Ali, S. S.; Abbasi, A. A.; Mohammad, A.; Wei, D. Higher Infectivity of the SARSlJCoVlJ2 New Variants Is Associated with K417N/T, E484K, and N501Y Mutants: An Insight from Structural Data. Journal of Cellular Physiology 2021, jcp.30367.

(10) SARS-CoV-2 Variant Classifications and Definitions https://www.cdc.gov/coronavirus/2019-ncov/variants/variant-info.html (accessed 2021 -06 -14).

(11) Eijsink, V. G. H.; Bjørk, A.; Gåseidnes, S.; Sirevåg, R.; Synstad, B.; Burg, B. van den Vriend, G. Rational Engineering of Enzyme Stability. Journal of Biotechnology 2004, 113 (1–3), 105–120.

(12) Ó’Fágáin, C. Engineering Protein Stability. In Methods in molecular biology (Clifton, N.J.); Humana Press, 2011; Vol. 681, pp 103–136.

(13) Shorthouse, D.; Hall, M. W. J.; Hall, B. A. Computational Saturation Screen Reveals the Landscape of Mutations in Human Fumarate Hydratase. Journal of Chemical Information and Modeling 2021, 61 (4), 1970–1980.

(14) Lee, M.; Shorthouse, D.; Mahen, R.; Hall, B. A.; Venkitaraman, A. R. Cancer-Causing BRCA2 Missense Mutations Disrupt an Intracellular Protein Assembly Mechanism to Disable Genome Maintenance. Nucleic Acids Research 2021, 49 (10), 5588–5604.

(15) Fowler, J. C.; King, C.; Bryant, C.; Hall, M. W. J.; Sood, R.; Ong, S. H.; Earp, E.; Fernandez-Antoran, D.; Koeppel, J.; Dentro, S. C.; Shorthouse, D.; Durrani, A.; Fife, K.; Rytina, E.; Milne, D.; Roshan, A.; Mahububani, K.; Saeb-Parsy, K.; Hall, B. A.; Gerstung, M.; Jones, P.H. Selection of Oncogenic Mutant Clones in Normal Human Skin Varies with Body Site. Cancer Discovery 2021, 11 (2), 340–361.

(16) Teng, S.; Sobitan, A.; Rhoades, R.; Liu, D.; Tang, Q. Systemic Effects of Missense Mutations on SARS-CoV-2 Spike Glycoprotein Stability and Receptor-Binding Affinity. Briefings in Bioinformatics 2021, 22 (2), 1239–1253.

(17) Smaoui, M. R.; Yahyaoui, H. Unraveling the Stability Landscape of Mutations in the SARS-CoV-2 Receptor-Binding Domain. Scientific Reports 2021, 11 (1), 9166.

(18) Walls, A. C.; Park, Y.-J.; Tortorici, M. A.; Wall, A.; McGuire, A. T.; Veesler, D. Structure, Function, and Antigenicity of the SARS-CoV-2 Spike Glycoprotein. Cell 2020, 181 (2), 281–292.e6.

(19) Public Health England. Investigation of novel SARS-COV-2 variant Variant of Concern 202012/01 https://assets.publishing.service.gov.uk/government/uploads/system/uploads/attachment_data/file/959438/Technical_Briefing_VOC_SH_NJL2_SH2.pdf (accessed 2021 -06 -14).

(20) Public Health England. Investigation of novel SARS-CoV-2 variant (202012/01): Technical briefing 2 https://assets.publishing.service.gov.uk/government/uploads/system/uploads/attachment_data/file/959361/Technical_Briefing_VOC202012-2_Briefing_2.pdf (accessed 2021 -06 -14).

(21) Ramanathan, M.; Ferguson, I. D.; Miao, W.; Khavari, P. A. SARS-CoV-2 B.1.1.7 and B.1.351 Spike Variants Bind Human ACE2 with Increased Affinity. The Lancet Infectious Diseases 2021, 0 (0).

(22) Schymkowitz, J.; Borg, J.; Stricher, F.; Nys, R.; Rousseau, F.; Serrano, L. The FoldX Web Server: An Online Force Field. Nucleic Acids Research 2005, 33 (Web Server), W382– W388.

