## Supplementary text for "SARS-CoV-2 Variants are Selecting for Spike Protein Mutations that Increase Protein Stability"

### Experimental procedures:

#### ΔΔG Calculation:

To study the mutational landscape of the SARS-CoV-2 spike protein from PDBID 6VXX[1], the structure was initially relaxed and repaired using the RepairPDB command in Foldx4[2] as follows:

```
$foldx --command=RepairPDB --pdb=5upp.pdb --ionStrength=0.05
--pH=7 --vdwDesign=2
```

RepairPDB was repeated on the structure five times to minimize its energy. The relaxed structure was then used to calculate the ΔΔG. PositionScan was run on each residue in the protein structure sequentially using the following command:

```
$foldx --command=PositionScan --pdb=6vxx_repaired.pdb
--ionStrength=0.05 --pH=7 --vdwDesign=2 --pdbHydrogens=false
--positions=100
```

To run PositionScan on the 100$^{th}$ residue.

#### Mutations:

Mutations in SARS-CoV-2 variants were obtained from CoVariants[3] (https://covariants.org/)

.

#### Expected mutational ΔΔG:

To calculate the expected mutational ΔΔG for a variant (Supplementary Figure 1), 1,000,000 samples of the same number of mutations in the variant were taken from the structure. For each sample the ΔΔG was calculated and the median of the distribution taken as the expected value. The value observed for the variant was removed from the expected to generate the ΔΔG difference.

#### Mutational ΔΔG combinations:

To calculate the ΔΔG for combinations of mutations in each variant, every possible combination of mutations in each variant was calculated. Each combination was then generated 15 times and  average ΔΔG calculated using the Foldx BuildModel command:

```
$foldx --command=BuildModel --pdb=6vxx_repaired.pdb --mutant-
file=mutantfile.txt --numberOfRuns=15 --pH=7 --vdwDesign=2 --
ionStrength=0.05
```

Where mutant-file.txt is a file containing the mutational combination to be modelled separated by a comma. For example, to model mutations L452R, D614G, and D950N in the Delta variant the file would contain:

LA452R,DA614G,DA950N;

**Supplementary Figures:**

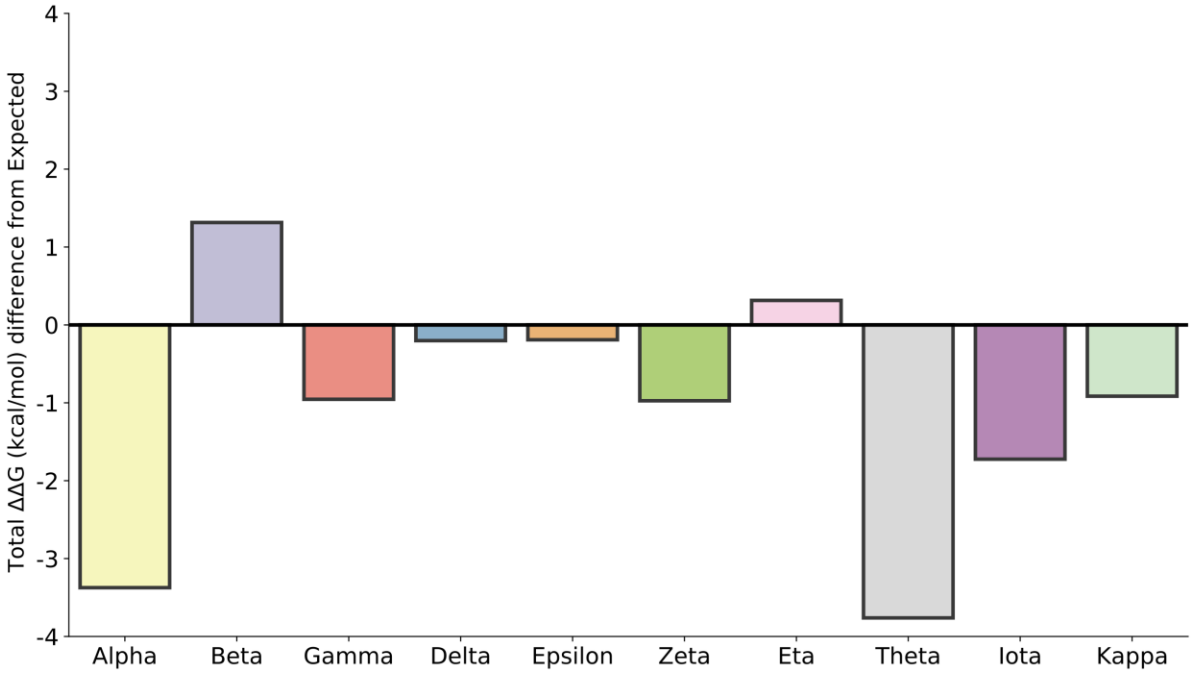


**Figure S1:** Difference between median expected ΔΔG for each variant and observed ΔΔG (Kcal/mol)

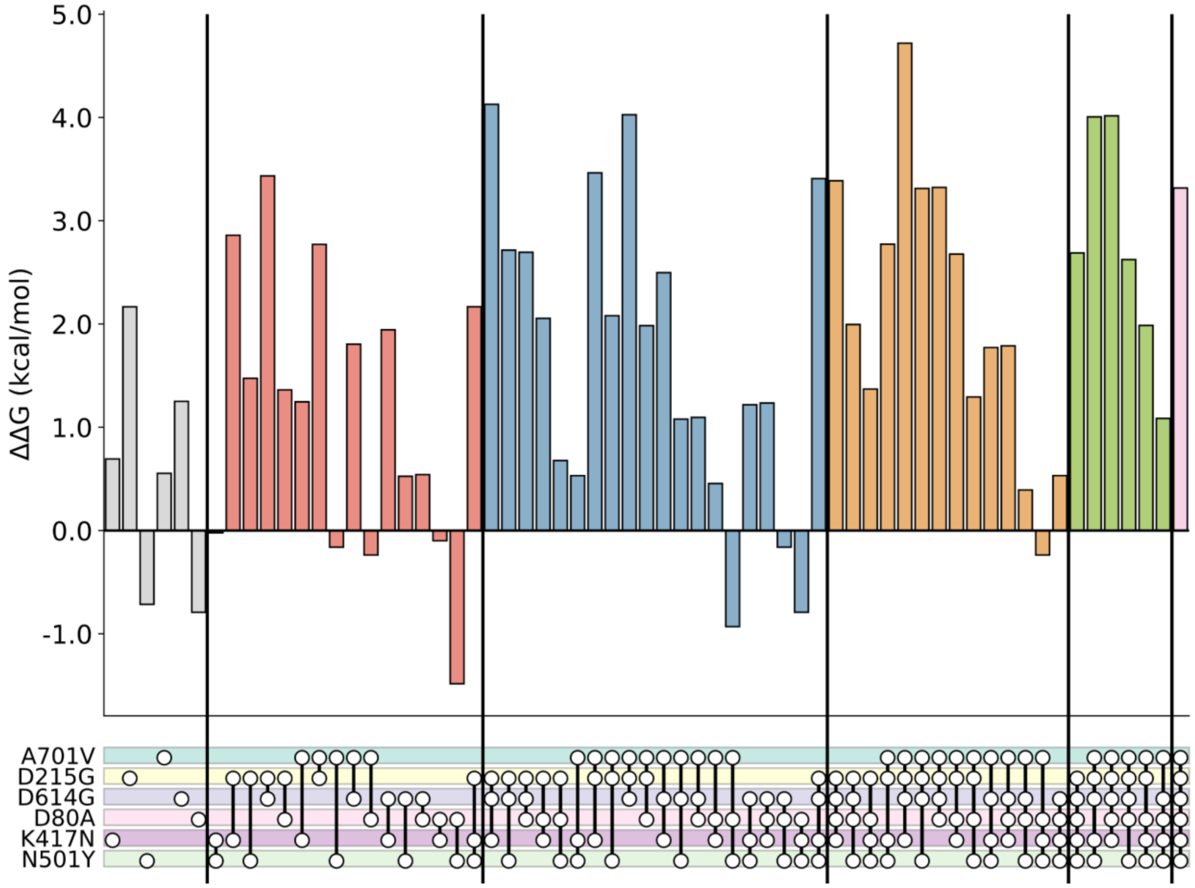


**Figure S2:** Upset plot for mutation combinations in SARS-CoV-2 Beta variant.

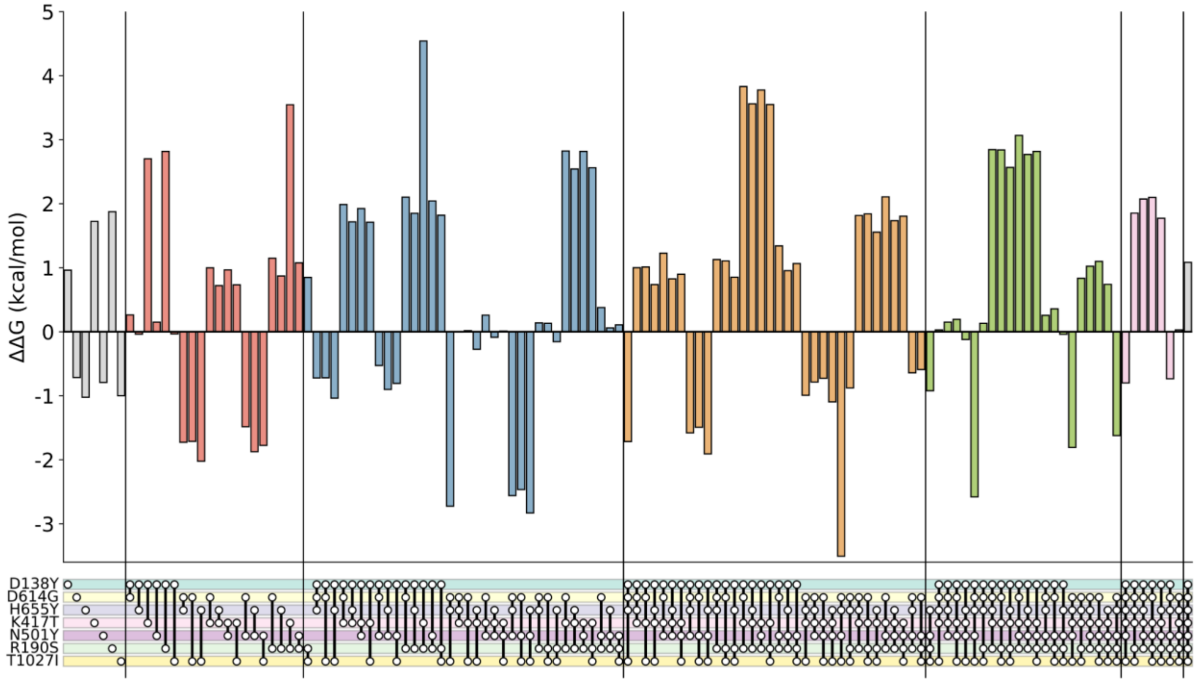


**Figure S3:** Upset plot for mutation combinations in SARS-CoV-2 Gamma variant.

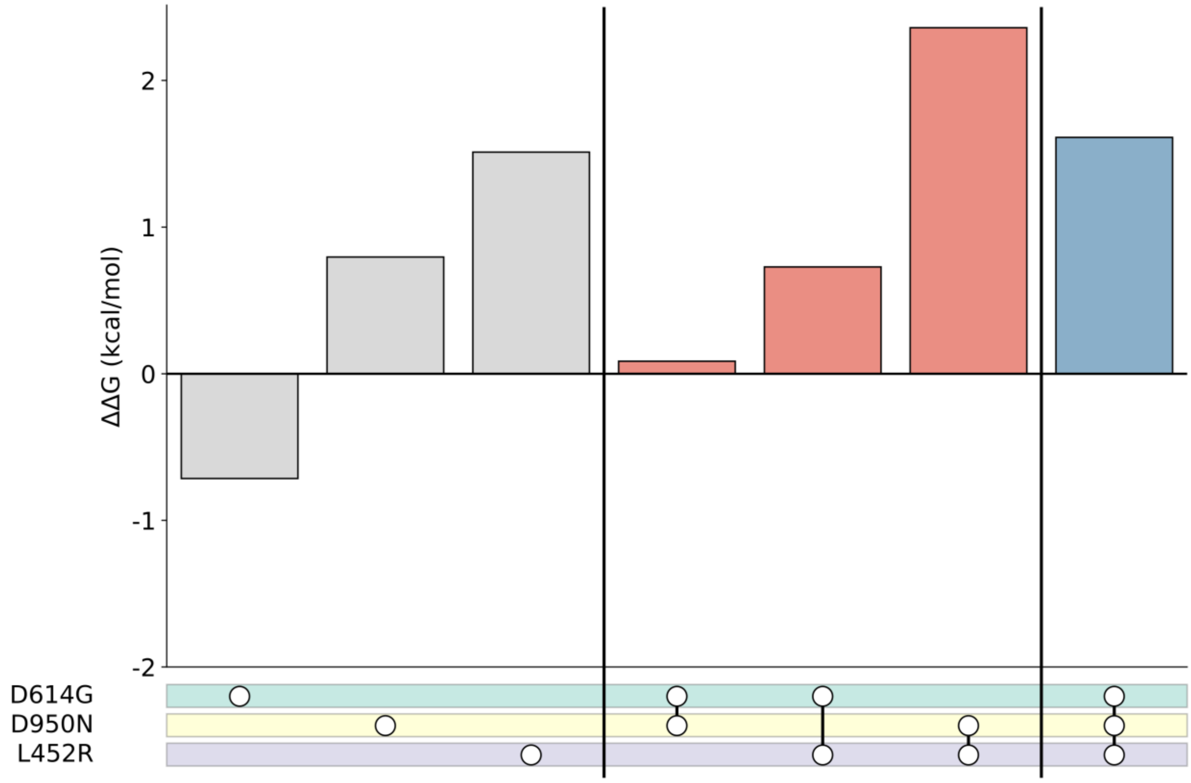


**Figure S4:** Upset plot for mutation combinations in SARS-CoV-2 Delta variant.

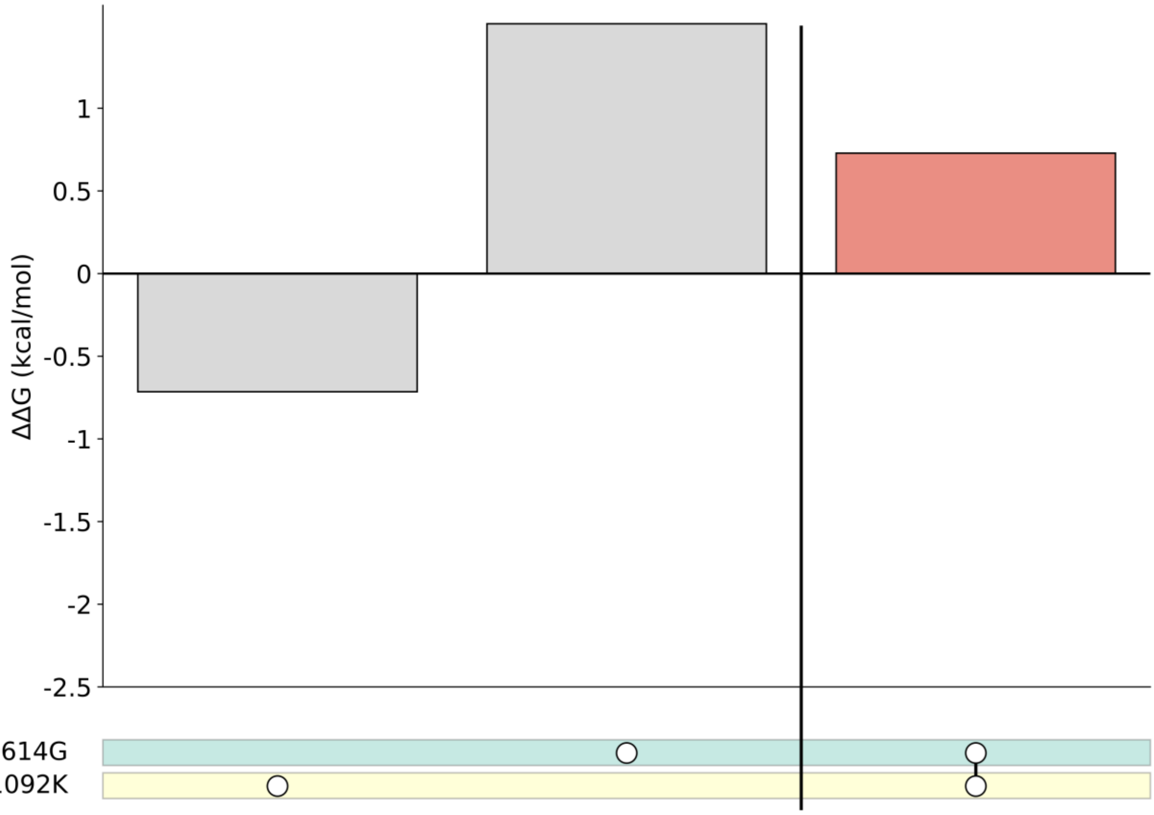


**Figure S5:** Upset plot for mutation combinations in SARS-CoV-2 Epsilon variant.

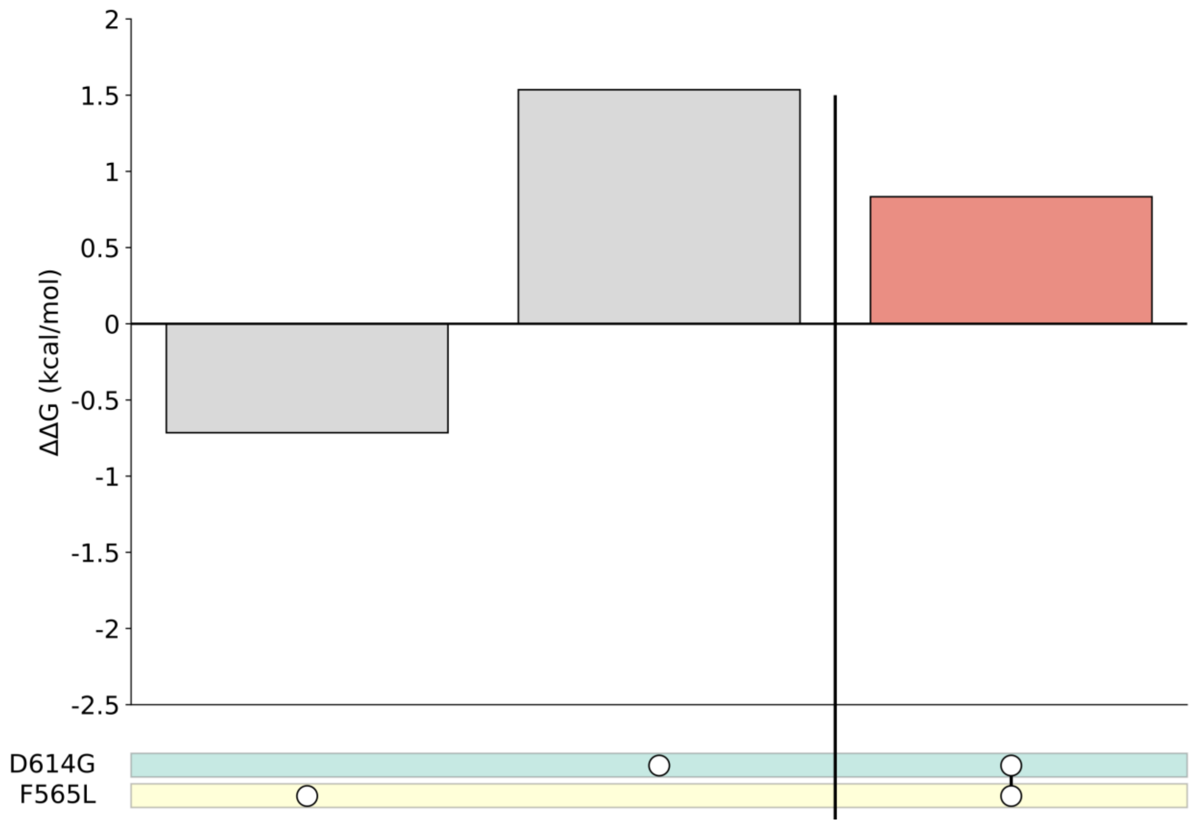


**Figure S6:** Upset plot for mutation combinations in SARS-CoV-2 Zeta variant.

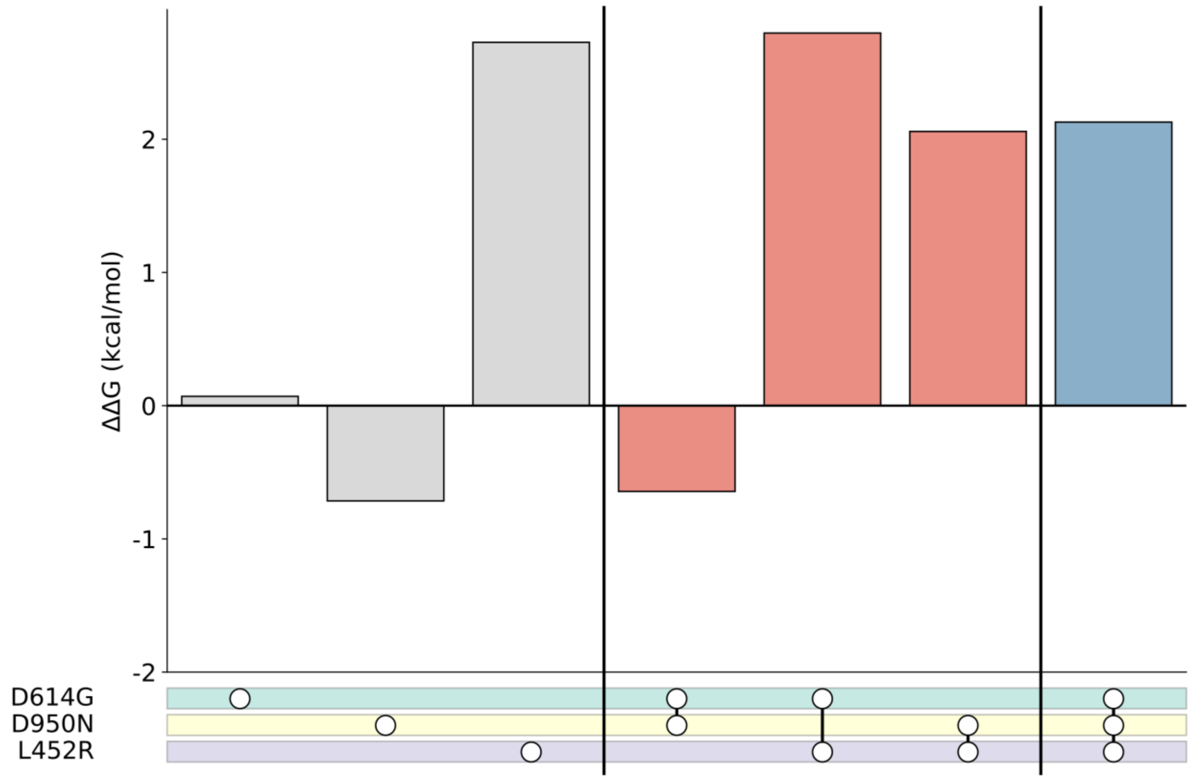


**Figure S7:** Upset plot for mutation combinations in SARS-CoV-2 Eta variant.

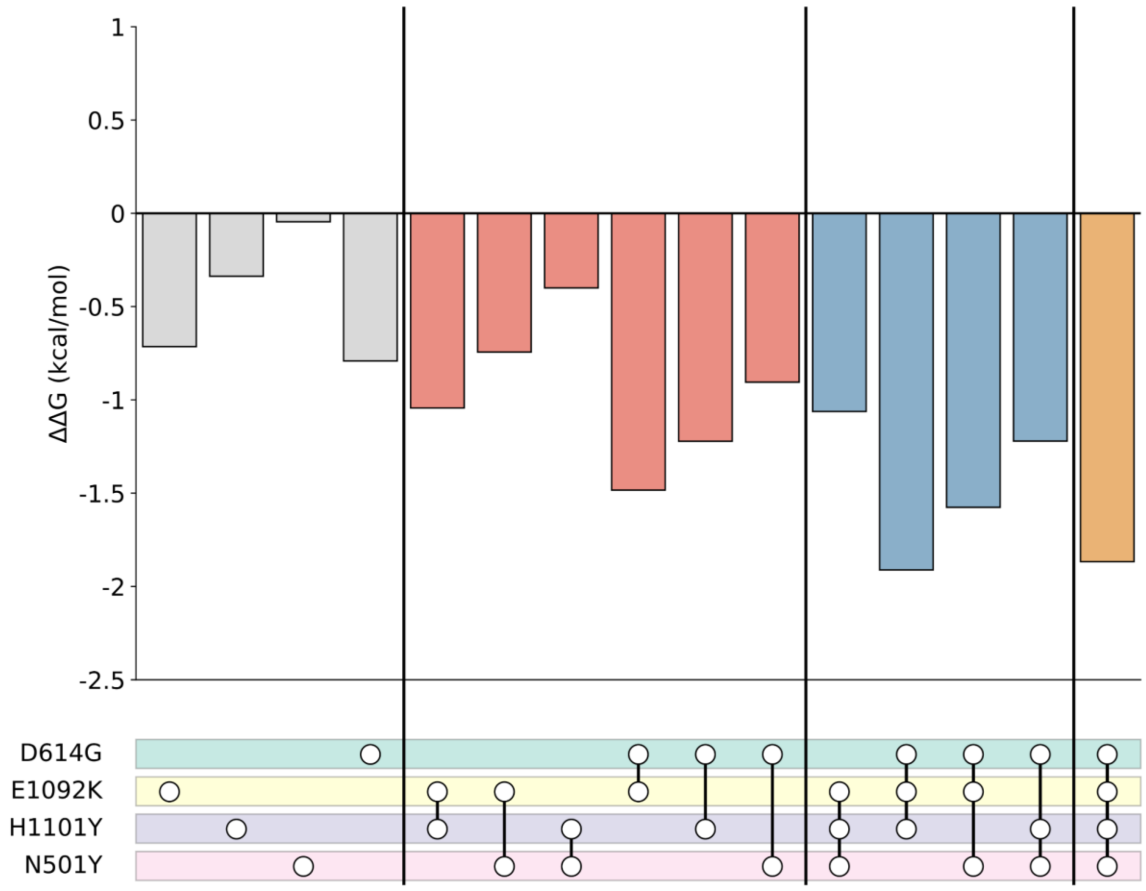

**Figure S8:** Upset plot for mutation combinations in SARS-CoV-2 Theta variant.

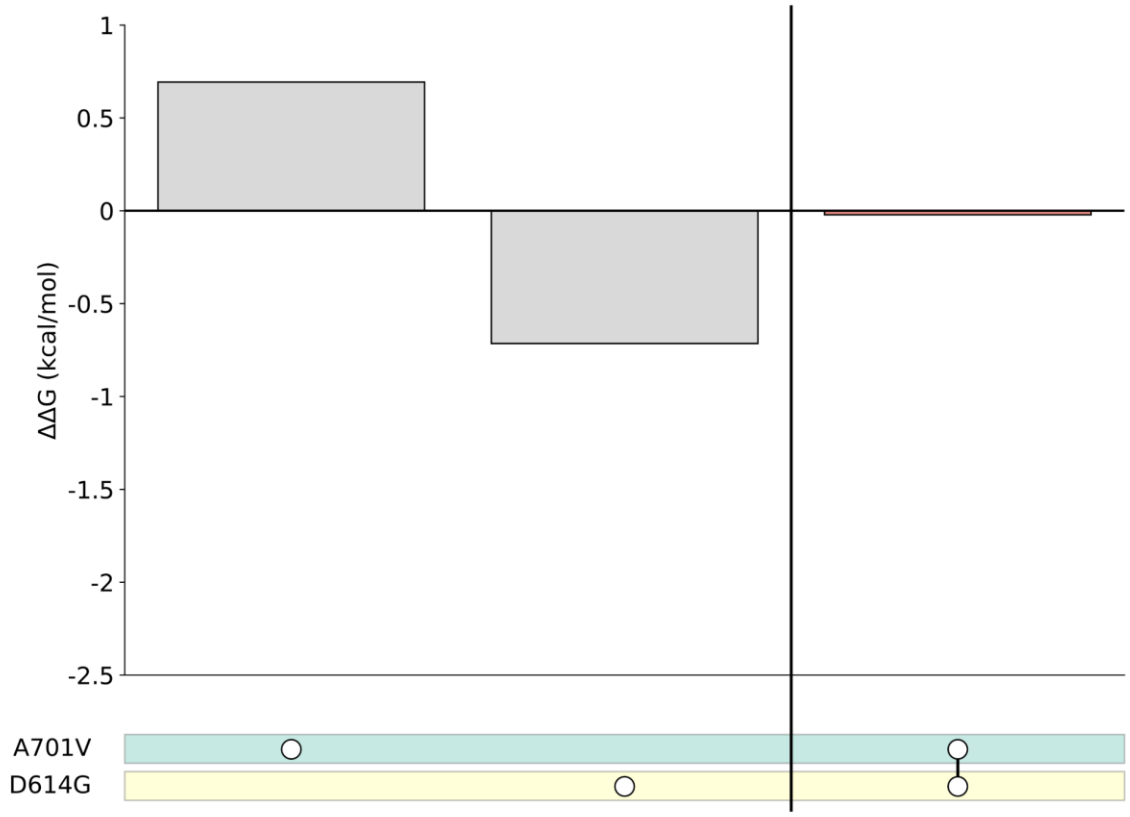

**Figure S9:** Upset plot for mutation combinations in SARS-CoV-2 Iota variant.

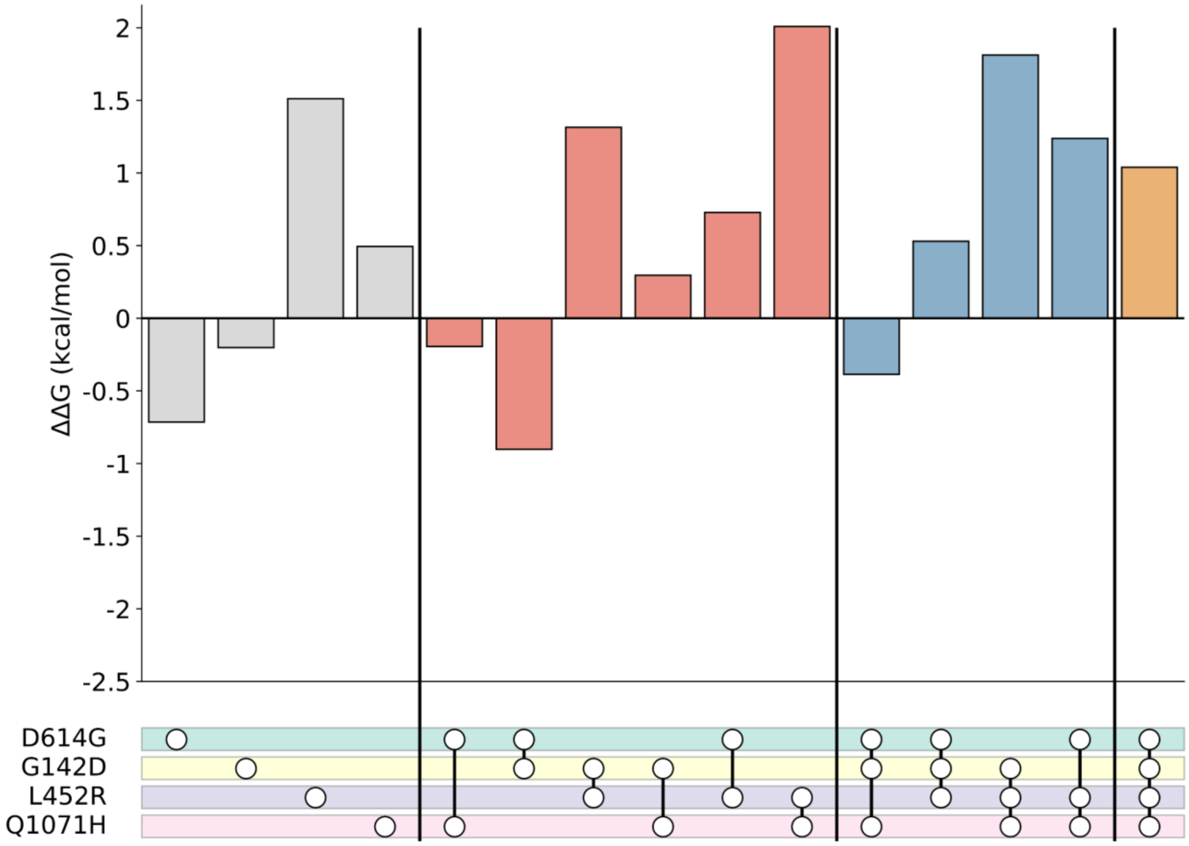


**Figure S10:** Upset plot for mutation combinations in SARS-CoV-2 Kappa variant.

**References:**


(1) Walls, A. C.; Park, Y.-J.; Tortorici, M. A.; Wall, A.; McGuire, A. T.; Veesler, D. Structure, Function, and Antigenicity of the SARS-CoV-2 Spike Glycoprotein. *Cell* **2020**, *181* (2), 281-292.e6. https://doi.org/10.1016/j.cell.2020.02.058.

(2) Schymkowitz, J.; Borg, J.; Stricher, F.; Nys, R.; Rousseau, F.; Serrano, L. The FoldX Web Server: An Online Force Field. *Nucleic Acids Research* **2005**, *33* (Web Server), W382–W388. https://doi.org/10.1093/nar/gki387.

(3) Emma B. Hodcroft. CoVariants: SARS-CoV-2 Mutations and Variants of Interest https://covariants.org/ (accessed 2021 -06 -15).
